# ^1^Arche: An Advanced Flexible Tool for High-throuput Annotation of Functions on Microbial Contigs

**DOI:** 10.1101/2022.11.28.518280

**Authors:** Daniel Gonzalo Alonso-Reyes, Virginia Helena Albarracin

## Abstract

The growing amount of genomic data has prompted a need for less demanding and user friendly functional annotators. At the present, it’s hard to find a pipeline for the annotation of multiple functional data, such as both enzyme commission numbers (E.C.) and orthologous identifiers (KEGG and eggNOG), protein names, gene names, alternative names, and descriptions. In this work, we provide a new solution which combines different algorithms (BLAST, DIAMOND, HMMER3) and databases (UniprotKB, KOfam, NCBIFAMs, TIGRFAMs, and PFAM), and also overcome data download challenges. The software framework, Arche, herein demonstrated competitive results over *Escherichia coli* K-12 genome, the metagenome-assembled genome of *Ferrovum myxofaciens* S2.4, and a freshwater metagenome when compared to other annotators. Finally, Arche provides an analysis pipeline that can accommodate advanced tools in a unique order, creating several advantages regarding to other commonly used annotators.

## I. Introduction

**A**crucial step in assessing the metabolic role of a microorganism is functional gene prediction and metabolic reconstruction [1]. The functional data “annotated” by means of bioinformatic tools can be used to infer metabolic differences with close relatives, characterize novel potential metabolisms, or detect the presence of antibiotic-resistance genes or toxins, among other genes of interest.[2], [3].

Genome annotation is the process of identifying and labeling all the relevant features in a genome sequence [4]. This should include coordinates of coding regions and their predicted products, but it may also be desirable to identify non-coding RNAs, signal peptides, etc. The genome annotation starts with gene identification or gene calling (structural annotation), which can be automatically performed using several tools, including Prodigal and GeneMark among others[5], [6], [7]. The next annotation step relies on the use of reference protein databases (DBs) to assign functions to query (unknown) protein sequences based on homology or orthologues searches (functional annotation) [8], [9]. Some of the DBs used are comprehensive, including protein sequences derived from complete and partial genomes and are usually updated periodically; widely used examples include the Uniprot[10], RefSeq[11], InterPro[12], Pfam[13] and SEED[14] DBs. On the other hand, specialized DBs aim at curating the entries to include only protein sequences related to specific functions or protein families of interest, e.g., CARD for antibiotic resistance proteins [15]. The search tools used against the DBs are variable, and in some cases optional, being BLAST[16], DIAMOND [17], SWORD [18] or HMMER [19] the most used.

The current annotation pipelines vary in complexity, comprehensiveness, speed, scalability, requirements and outputs. For example, the NCBI Prokaryotic Genome Annotation Pipeline (PGAP)[20], Prokka[21], RAST[22], Bakta[23], and DRAM [24] start from genomic contigs and predict genes and proteins, tRNAs, rRNAs, and perform functional annotation of the predicted proteins. Others such as InterProScan[25], eggNOG-Mapper[8], [26] and MicrobeAnnotator[27] start from already predicted protein sequences. The RAST service [22] requires an account and email, with a turn-around time measured in less than a day. It’s almost the same with PGAP, a web server that provides annotation results in several days [22]. On the other hand, Prokka, MicrobeAnnotator, DRAM, and eggNOG-Mapper can be run in a desktop computer but they have limitations. Prokka relies on a very short database (DB) and generates a constrained number of annotated proteins. At the opposite, MicrobeAnnotator in standard mode demands the download of a huge DB (250-300 GB) which requires a lot of disk space and fast internet connection. eggNOG-Mapper is circumscribed to the eggNOG DB[28] and fails to combine different search algorithms (e.g. BLAST and HMMER) in the same run. Finally, Bakta is intended only for bacterial genomes, and DRAM is designed to be run on high-performance computers.

As a result of the pros and cons of the above-mentioned tools, it is necessary to develop an annotation pipeline that simultaneously accounts for the following: 1) resolve trait prediction from vast amounts of genomic content; 2) combine diverse databases and algorithms in the same run; 3) be able to launch from a desktop or laptop; 4) be easy to download. Additionally, the software should produce highly informative annotation tables for constructing metabolic pathways, comparing orthologues, and screening single traits. We present Arche, a bash-based command-line tool designed for the coordinated and optimized annotation of prokaryotic contigs using a suite of existing software tools. With Arche, several genomes can be annotated simultaneously or the annotation of an individual genome or metagenome can be speed up by taking advantage of multiple processing cores.

## II. Implementation

### A. Input

Arche expects previously assembled genomic or metagenomic DNA sequences in FASTA format. This sequence file is the only mandatory parameter to the software. The tool is not usable if the user has only proteins sequences as input.

### B. Annotation Process

Before functional annotation can be made, coding sequences (CDS), ribosomal RNA genes (rRNAs), and transfer RNA genes (tRNAs) must be discovered inside the contigs. Arche relies on external prediction tools to identify those features within the contigs (Fig. 1). These tools are:

**Fig. 1.**
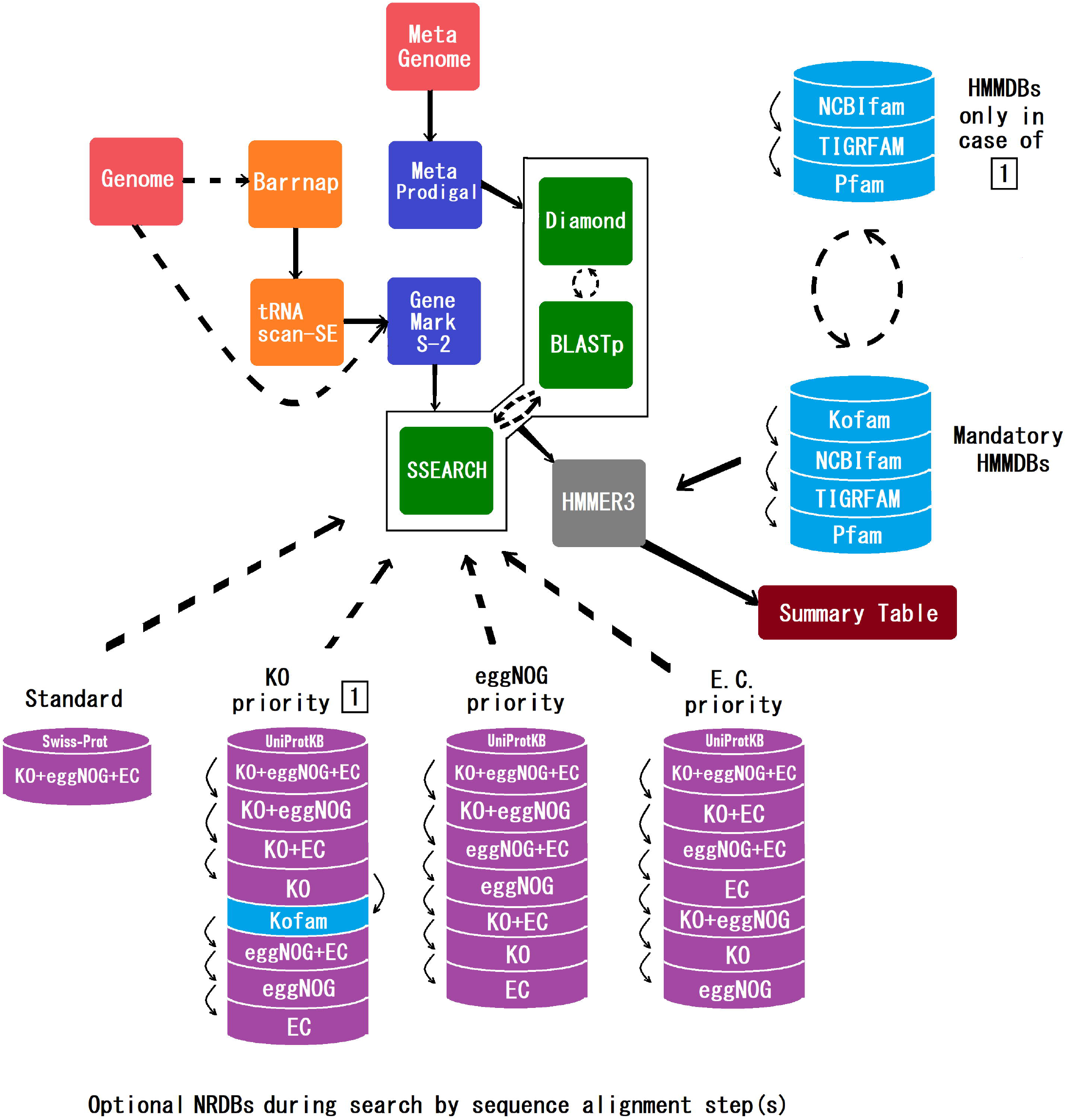
Arche workflow. The RNA prediction stage (orange boxes) is optional (broken lines) and only applied to genomes. GeneMarkS-2 and MetaProdigal are used to perform protein prediction (blue boxes) from genomes and metagenomes respectively. During the first search stage (green boxes), BLASTp, DIAMOND, or SSEARCH can be used alternatively to align the input proteins over the four alternative NRBDs (purple containers). Each of those NRBDs are split in several clusters of sequences which have functional information associated (KO identifiers, eggNOG identifiers or Enzyme Commission numbers). Except the “standard” NRBD, each one prioritizes the annotation of a particular functional identifier. During the second search stage (grey box) the remnant unannotated proteins serve as input for HMMER which scan over several HMMBDs (cyan containers). In “KO priority” the NRBD is interrupted by KOfam DB (cyan container) after the fourth cluster, which means that all the input sequences unannotated by the previous clusters would be scanned on KOfam DB using HMMER. After this, the workflow will resume the alignment with the remaining clusters of the “KO priority” NRBD. The final output of the workflow is a summary table with biological information associated to the matches.

(1) barrnap (https://github.com/tseemann/barrnap) for rRNAs.

(2) tRNAscan-SE 2.0 [29] for tRNAs.

(3) GeneMarkS-2 [30] for coding sequences (CDS) in genomes.

(4) MetaProdigal[31] for coding sequences (CDS) in metagenomes.

Once the coordinates of candidate gene sequences have been identified, a description is required of the putative gene product. The traditional method of predicting a gene’s function is to compare its sequence against a set of sequences (database) with already known functions. Then, the CDS is labeled with the name and function of the closest significant match. Arche uses this method, but in a hierarchical manner similar to Prokka[21]. Depending of the option provided by the user, it starts making comparisons by heuristic (e.g. BLAST) or Smith-Waterman (SSEARCH) algorithms over in-house made databases, then uses a hmm algorithm over databases of protein families, and finally the same hmm algorithm over a domain-specific database. In each step, those sequences that didn’t matched any homolog in the DB pass to the next step. In summary Arche queries the genomic CDS over the following series of DBs:

1. A set of sequences picked from the UniProtKB DB and clustered in new redesigned DBs (NRDBs). These sequences have an ID number and an associated entry linked to functional information (description, name, alternative name, E.C. number, etc). To create the NRDBs, we first filtered the IDs of interest using the options “AND, OR, NOT” from the advanced search of the Uniprot website (http://www.uniprot.org/). This filtering was made with the goal of cluster the IDs having associated Kegg Orthologues (KO[9]), eggNOG orthologues (eggNOG[28]), or Enzyme Commission numbers (E.C.[32]), and exclude those IDs lacking these identifiers. Once the clusters were defined, the IDs were used to download the fasta sequences and build the NRDBs. No de-replication process was performed over the sequences. Four optional NRDBs are provided by Arche; three of them have the ability to bias the annotation of one of three functional codes— KO, eggNOG, or E.C. The fourth NRDB is built in the same way than the others but from the Swiss-Prot DB, and it enables the annotation of any of the three aforementioned functional codes (no bias). The search over the NRDBs is made by a heuristic or Smith-Waterman algorithm with a default e-value threshold of 10^−8^. All NRDBs were built with domain specificity, i.e. with bacterial or archaeal sequences only. In case of the annotation of metagenomic contigs, the program will search a combined NRDB made with both bacterial and archaeal sequences.
2. A series of HMM profile DBs (HMMDBs) in this order: a prokaryotic version of KOfam[33], NCBIFAMs [34], TIGRFAMs [35] and Pfam[13]. The hmm algorithm of HMMER3 [19] is used for the search. By default, gathering score thresholds (implemented with the --cut_ga option) are used during the search over the HMMDBs except KOfam. In the case of KOfam search, the pre-computed adaptive score thresholds[33] were used (incorporated in the hmm file and implemented with the --cut_ga option).
3. If no matches can be found, the CDS is labelled as ‘hypothetical protein’.

Table I describes the DBs used, their sources, and respective modifications.

**Table I.**
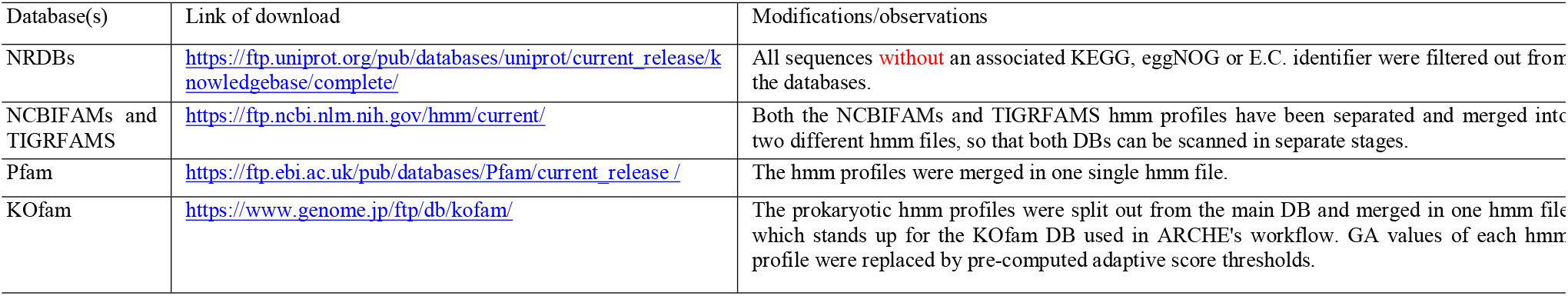
Description of DBs used by Arche, their sources, and respective modifications.

To provide greater flexibility to the user in the querying over the NRDBs clusters, Arche supports the use of three different popular search tools, i.e., BLASTp[16], DIAMOND[17] and SSEARCH[36]. The user can also modify the filtering thresholds used to select the best match found in the DBs (i.e., e-value, or query cover). Finally, multiprocessing is also supported with the - t (threads) option.

There are four modes of running Arche, “standard”, “kegg”, “eggnog”, and “ec”. The “standard” mode uses a reduced NRDB (based on Swiss-Prot DB) that annotates KO, eggNOG, or E.C. codes, before starting using HMMDBs. The remaining modes perform almost the same pattern as the “standard” scanning but replaces the Swiss-Prot DB with an NRDB made from sequences from UniprotKB. These alternative modes bias the annotation to E.C., KO or eggNOG codes, depending on the option provided by the user, “kegg”, “eggnog”, or “ec”, respectively. Also the NRDBs assigned to these modes are subdivided into seven sequence clusters (Fig. 1, purple discs). At the top of the scanning are those clusters of sequences that have the functional codes prioritized (e.g. KO), and at the bottom are the remaining clusters. The scanning will proceed through all seven clusters before starting with HMMDBs. For example, if the mode is “ec”, the clusters will be searched in this order: KO + eggNOG + E.C. (cluster 1), KO + EC (cluster 2), eggNOG + EC (cluster 3), EC (cluster 4), KO + eggNOG (cluster 5), KO (cluster 6), and eggNOG (cluster 7), after which HMMDBs will begin to run. If the mode is “kegg”, then the search will be as follows: KO + eggNOG + E.C. (cluster 1), KO + eggNOG (cluster 2), KO + EC (cluster 3), KO (cluster 4), KOfam(HMMDB), eggNOG + E.C (cluster 5), eggNOG (cluster 6), and EC (cluster 7). In the last example it’s noticeable that KOfam scanning begins after the last cluster that contains KO, i.e. cluster 4 (Fig. 1), thus truncating cluster scanning. Once KOfam search is over, truncated cluster scanning will be resumed, and then HMMDBs will run (except KOfam). This procedure is intended to maximize the annotation of KEGG IDs (KO).

At the end of the annotation, query sequences are labeled with the database-specified code of the matching sequence. This code will be used by the pipeline to retrieve the metadata from each of mapping tables implemented in Arche. This final table will summarize all the functional information (names, KO identifiers, E.C. numbers, etc.) provided by the different DBs queried. However, some data is not linked directly. For example, the Uniprot-associated KO identifiers were obtained through the conversion of the KEGG identifiers to KO identifiers via script. This procedure involves the download of the KEGG entry and the extraction of the KO ID (Fig. 2). Moreover, the E.C. numbers associated to the Pfam DB were obtained from the ECDomainMiner approach [37]. At the end, all annotated CDS will be grouped in a single table per genome, which includes all metadata associated with each best match.

**Fig. 2.**
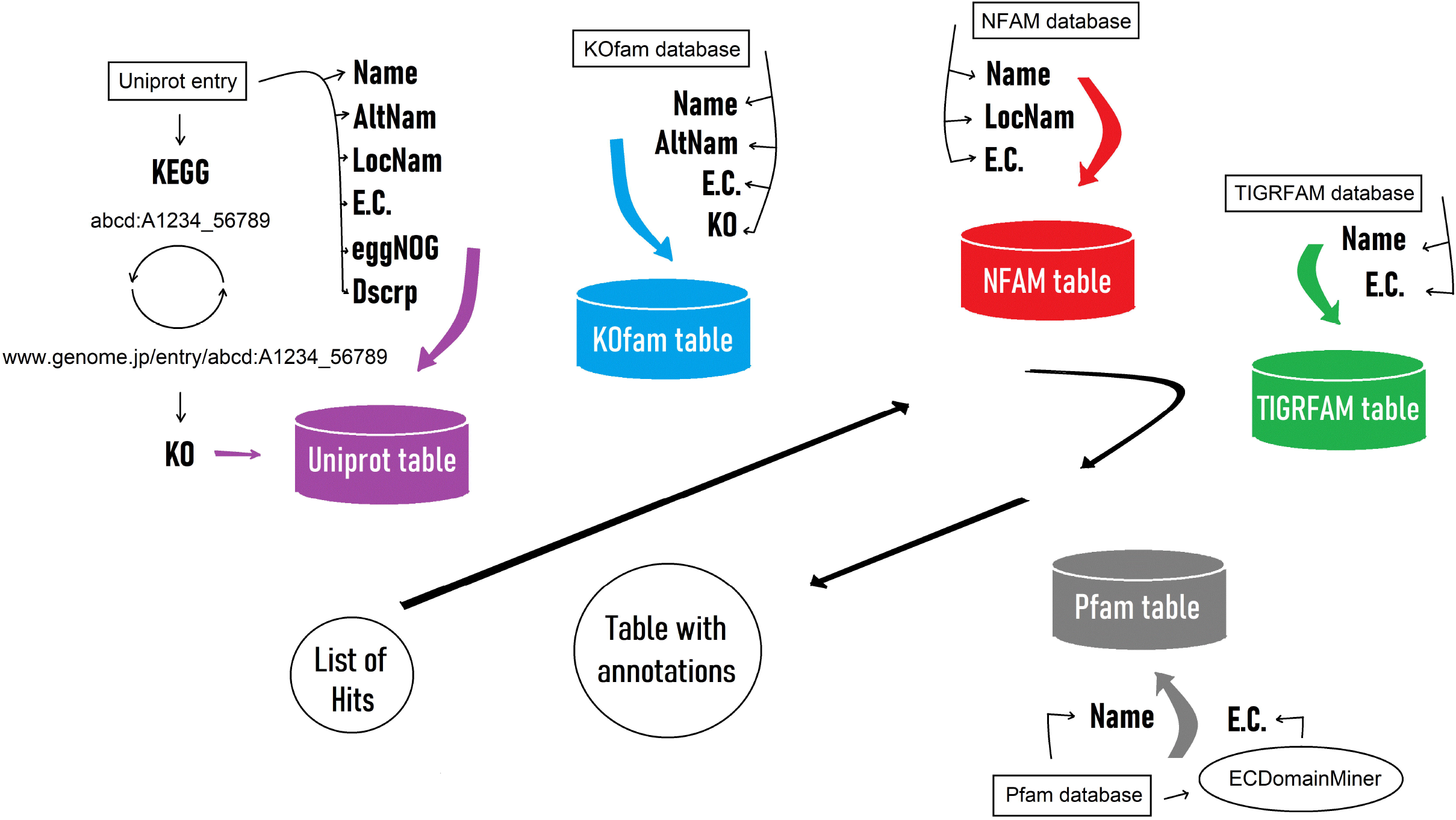
Picture showing the annotation process. Once a group of hits from the search step(s) is obtained, their respective identifications are matched to the functional data saved in tables (colored containers). Each table gathers all the useful information provided by the original source (Uniprot, KOfam, etc.). Finally, all the query identifiers, the query matches, and the corresponding related data will be resumed in one single table.

### C. Output

Arche produces a single main output folder per run, which contains several files described in Table II. Snippets from files []_omic_table.tbl, []_omic_table.tsv, and arche_report are showed by Supplementary Figs. 1-3.

**Table II.**
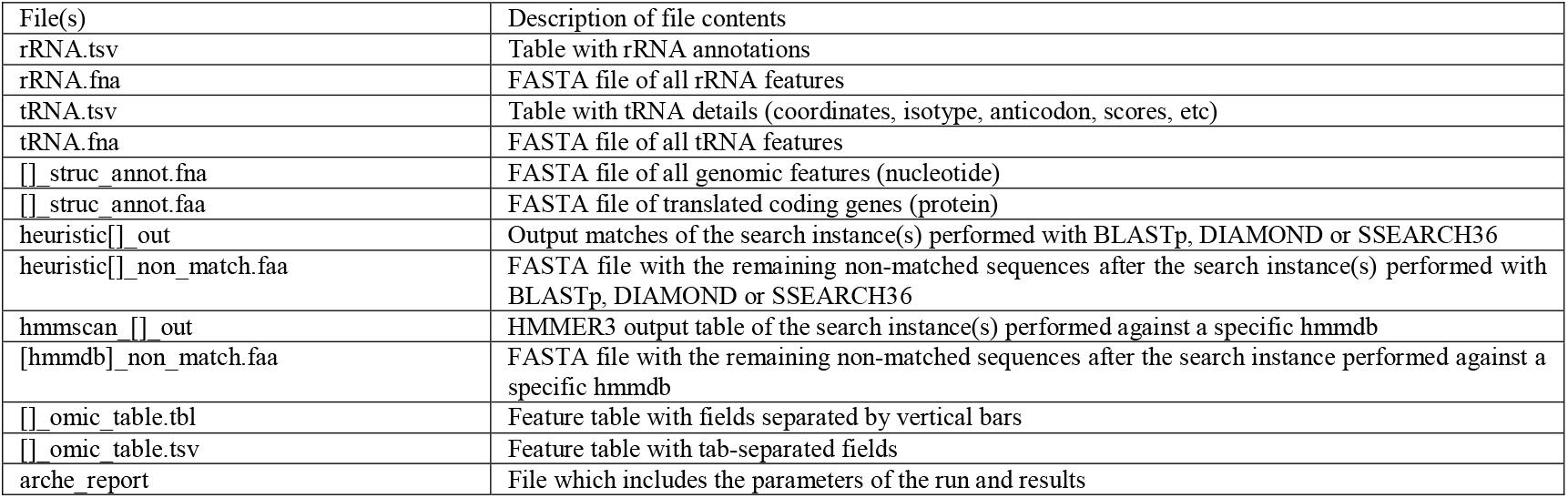
Description of Arche output files.

## III. Results

### A. Computing Requirements of Arche compared to other tools

We compared Arche to other popular genomic annotation pipelines, including Prokka[21], RAST[22], eggNOG-Mapper[26], Bakta[23], and MicrobeAnnotator light mode[27].

The first comparison was in terms of download size of the required DBs to perform an annotation. These DBs include FASTA protein entries, HMM profiles, and mapping files depending on the tool. Table III shows that Bakta (39,4 Gb) and Arche (13 Gb) required downloads are bigger than the other tools. Arche’s download size is mostly driven by the inclusion of the NRDBs, four different HMMDBs, and their respective mapping tables. Those DBs are already formatted for all Arche’s modes and searching tools, and suitable for the annotation of both bacterial and archaeal genomes, and metagenomes. The Arche’s NRDBs include only those sequences associated with E.C./KO/eggNOG data while most of the sequences and map files used by the other programs have not such data associated.

**Table III.**
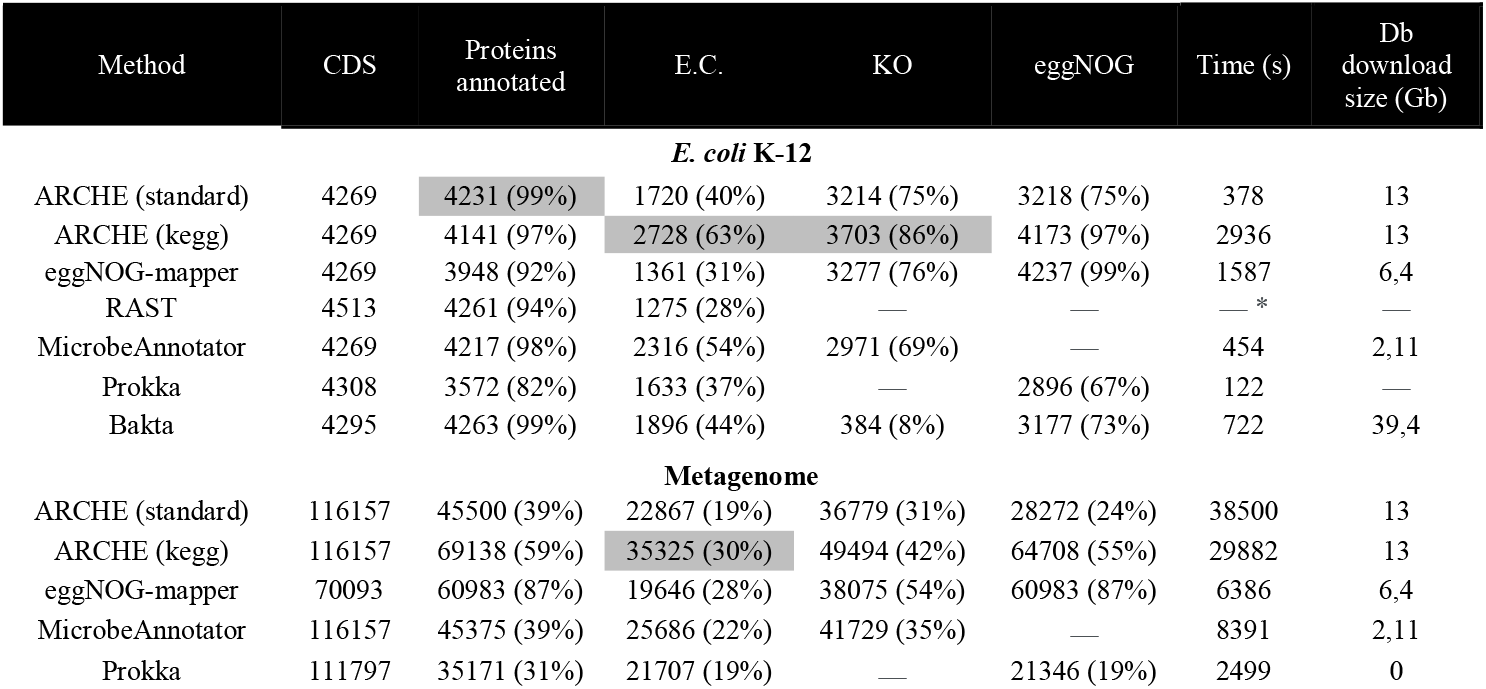
Comparison of the performance of several annotators, including Arche in the annotation of *Escherichia coli* K12 and a freshwater metagenome. The percentages of CDSs annotated are also indicated. Highlighted in grey are the metrics in which Arche is performing the best. *Depending of the job load of the server, RAST can take almost a day to return the results of the annotation, as it happened with the *E. coli* K-12 genome.

More important than the download size, we compared each tool’s annotation performance in terms of speed, and number of annotations. For this, we screened the genome of the well-known *Escherichia coli* K-12 strain (NCBI NC_000913.3), and a freshwater metagenome (GCA_900143125.1) with all the aforementioned tools including in the testing two modes of Arche: Arche “standard” mode (AS) and Arche “kegg” mode (AK). Of the tools tested, Prokka and AS were the fastest, taking 122 and 378 seconds to annotate the *E. coli* genome, respectively (Table III). These were followed by MicrobeAnnotator and Bakta which took 454 s and 722 s to annotate the genome. In our tests and depending on the server load, the web version of RASTtk can take almost a day. Finally, AK required significantly more time to fully annotate the genome (2936 s). The metagenome analysis led to a much slower behavior of Arche, in comparison to other tools. This is likely because a higher proportion of the protein sequences will reach the HMM analysis step, which is more time consuming than the previous steps. AS also will be slower than AK, because of its small NRDB which will cause relatively more of the protein sequences to reach the HMM analysis step.

### B. Annotation completeness of Arche compared to other tools

In general, all tools annotated 92-99% of the CDSs of the genome of *E. coli* K-12, except for Prokka, which annotated 82% (Table III). Overall, AS and AK had high percentages of annotation, with 97% and 99% of CDSs annotated respectively. MicrobeAnnotator also gave good results in terms of CDSs annotated, with the 98% being classified with a function. On the other hand, the range of annotation in the metagenome vary between 31% for Prokka, and 87% for eggNOG-mapper. However, the latter number is biased by the lower amount of CDS produced by eggNOG-mapper during the gene calling step. This becomes evident when compared to the other tools (∼70093 vs >110000 CDS).

Out of all the tools tested, RAST had the lowest number of functional identifiers, including Enzyme Commission numbers which accounts for the 28% of the CDS of the *E. coli*’s genome (Table III). On the other hand, eggNOG-mapper and Prokka included E.C. numbers in 31% and 37% of the CDS, respectively. At the top of performances are Bakta (44%), MicrobeAnnotator (54%), and AK (63%). In regards to KO and eggNOG ortholog DBs, RAST doesn’t provide such data in the annotation. The remaining methods consistently included KO and eggNOG identifiers to complement the text-based annotation descriptions. 69%, 75%, 76%, and 86% of the annotated CDS include KO identifiers according to MicrobeAnnotator, AS, eggnog-mapper and AK results respectively. Bakta led to a very small proportion of CDSs with a KO identifier. Egg-NOG identifiers are present in 67% of CDS according to Prokka annotation, 75% of CDS in AS results, 97% in AK results, and 99% of CDS in eggNOG-mapper annotations. In regards to the metagenome, Prokka led to the lowest number of functional identifiers, including E.C. and eggNOG IDs, both of them accounting for the 19% of the CDS (Table III). No KO IDs were provided in the Prokka’s file of annotations. On the other hand, both AK and eggNOG-mapper showed very good performances in functional ID annotation.

Although AK run times are longer than those for the other tools (Table III), the output is rich in annotated CDSs (Table III) with both names and functional codes. Moreover, if the DIAMOND option in AK is used, the runtimes can be reduced, without a significant loss in annotation performance (data not shown).

### B. Quality of the annotations

We evaluate the quality of the Arche’s annotation through a concordance analysis against other annotation tools in the genome of *E. coli* K-12, the MAG of *Ferrovum myxofaciens* S2.4 (NZ_CP053676.1), and a metagenome (Figs. 3-5, Supplementary Tables S1-S3). We observe that AS and AK have opposite trends in regards the amount of validated annotations (red bar plots) depending of the type of input. For a well referenced genome like that of *E. coli* K-12, AS performs relatively well in comparison to other tools, but much better than AK. On the other hand, AK had better results when applied to the metagenome. Moreover, in the MAG of *F. myxofaciens* S2.4 both AS and AK had an intermediate performance when compared to other tools. In regards to the annotation of additional biological information like gene names, KO, E.C. and, eggnog IDs (green, blue, black texture, and yellow barplots respectively), Arche had very good performances as a whole when compared to the other tools.

**Fig. 3.**
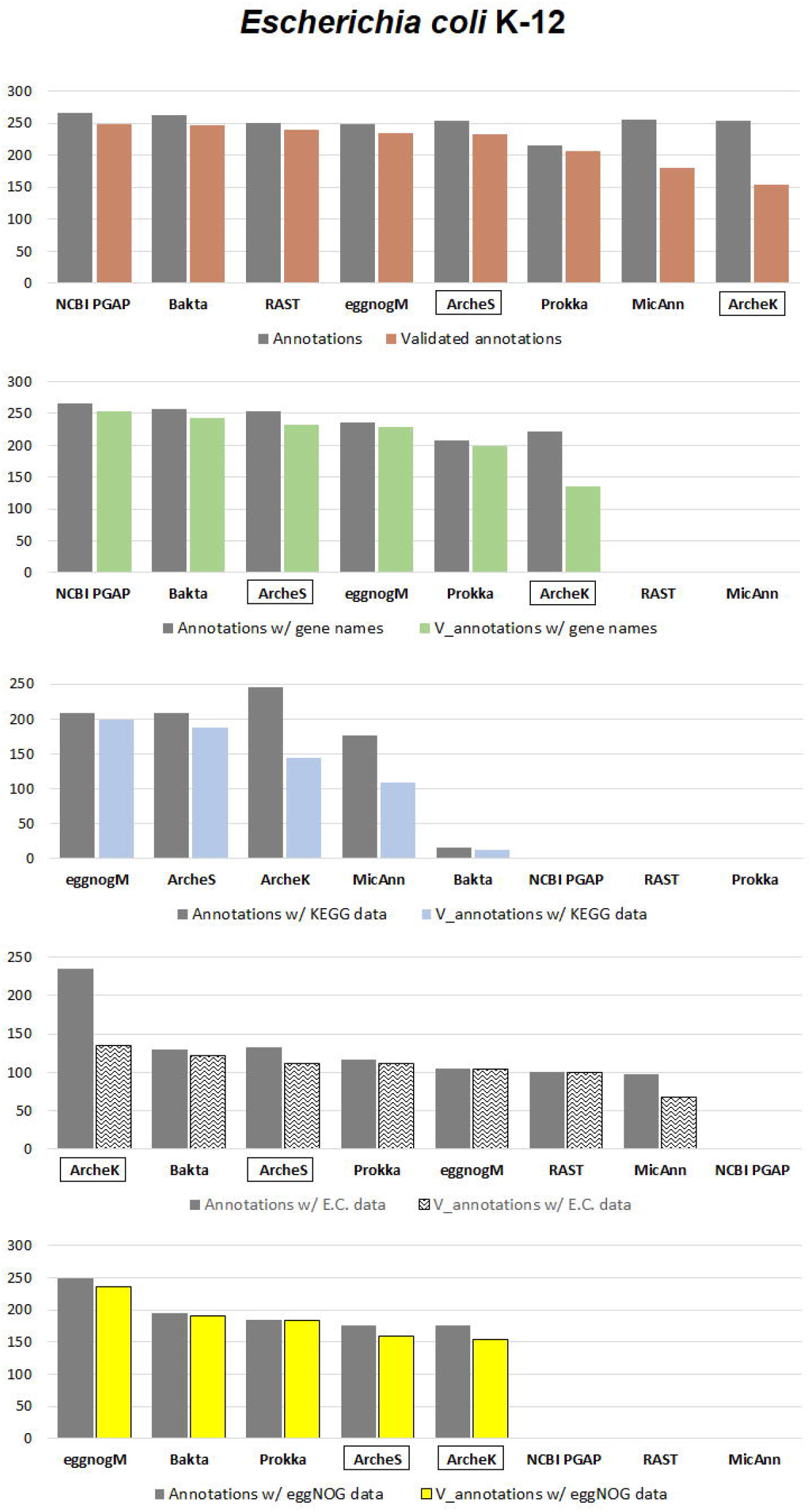
Bar plot that shows the performance of several annotation tools in the genome of *E. coli* K-12. Abbr. V_annotations = Validated annotations

## IV. Discussion

We provide a new software tool ‘‘Arche’’ a bash-based command-line tool for annotation of prokaryotic genomes. Arche presents a set of exclusive tools and procedures, while some of them are shared with the other annotators (Fig. 6). Theoretically, Arche have advantage in regards to the number of annotation steps, procedure options, or quality of some steps. For example, Arche is a complete genomic annotation pipeline since it can accept as input a genome (complete or fragmented) or even a metagenome. These contigs are going to be undergo structural annotation through the appropriate tools for the prediction of RNAs and proteins. MicrobeAnnotator lacks this option and depends on the user directly providing the already predicted protein fasta files. eggNOG-Mapper, on the other hand, does not map RNAs and performs protein gene calling only optionally. Regarding to the programs used for structural annotation, both Arche and Prokka use Barrnap (https://github.com/tseemann/barrnap) for the prediction of rRNAs. However, in the case of tRNAs, Arche exploits tRNAscan-SE 2.0[29], a more updated and improved program than ARAGORN[38] and tRNAscan-SE 1.0[22], [39] implemented in Prokka and RAST respectively. With regards to the protein gene calling in genomes, Arche implements the amazing GeneMarkS-2[30] tool which has been demonstrated to give better results than Prodigal[6] (used by Prokka and optionally by eggNOG-Mapper) and Glimmer[40] (used by RAST). Furthermore GeneMarkS-2’s output classifies coding regions in ‘native’ for species-specific genes and ‘atypical’ for harder-to-detect genes likely (horizontally transferred)[30]. In the case of metagenome annotation, both Arche and Prokka implement the MetaProdigal tool[31] which has been proven to be one of the best algorithms for metagenomic gene prediction. On the other hand, RAST has no option for metagenomic analysis at all.

When it comes to DBs used and search procedures, Arche benefits from summarizing other bigger DBs. The pipeline has four optional new redesigned DBs (NRDBs) made of picked sequences from the UniProtKB DB, and four different HMM profile DBs (HMMDBs) which are mandatory. All these DBs are run in a hierarchical strategy: first a search by sequence alignment (e. g. BLAST) is performed over a series DBs whose sequences have functional codes related, and then HMM searches (by using HMMER) are applied on the remaining sequences. This is based on the concept that HMM profile comparisons are more appropriate for detecting remote homologs[41].

Prokka uses the same logic although with less and shorter DBs. MicrobeAnnotator light mode reverses the procedure, launching an HMM search against the KOfam DB during the first step, and then applying a sequence alignment search against Swiss-Prot over the remaining sequences. eggNOG-Mapper fails to combine alignment search methods with HMM searches (they exist as alternative procedures), and RAST doesn’t have any HMM search at all. When we consider the search tools used during the sequence alignment step(s), Arche has some advances. It’s possible to choose between both BLASTp or DIAMOND as the main heuristic algorithms, or the SSEARCH implementation of the S-W algorithm. The options provided by MicrobeAnnotator are similar, although the SWORD tool replaces SSEARCH. However, in contrast to Arche, MicrobeAnnotator formats DBs according one of the three algorithms (e.g., BLAST), which would require a new DB download in case the user wants to run the annotation with another search tool (e.g., DIAMOND). eggNOG-Mapper has DIAMOND and MMseqs2 as alternatives, having MMseqs2 almost a similar sensitivity than BLAST, but still less at the end. Finally, both Prokka and RAST uses only BLAST in this step which makes them the least flexible of all the tools tested.

Another Arche’s advantage to the user is the retrieval of additional text-based information from the NRDBs and KOfam, like the locus name(s), or the alternative name(s) of a protein and text about its function whenever available (Fig. 2, 6). This gives the user an extra benefit in case of looking for particular features of interest using text search tools.

Arche provides a hierarchical workflow and incorporates state-of-the-art programs for both tRNA and protein prediction. The software demonstrated an innovative segmentation of steps towards a functional-oriented annotation of genes, and organized output of data within one tool. Overall, Arche provides an automated end-to-end solution for the entire annotation of prokaryotic contigs that is both flexible and suitable for extracting the kind of information that the user needs for a chosen downstream analysis. At present, only a small group of developed software produced the sufficient diversity and quantity of functional codes to allow adequate metabolic reconstruction. Some of these tools like DRAM demands high-performance computers; other require the download of huge datasets. For example, MicrobeAnnotator *standard* mode requires the download 200-300 Gb of DB only for a simple BLAST analysis. Some of the download blocks have a big size like the TrEMBL dat files (>150 Gb), thus requiring a fast and uninterrupted connection to be downloaded successfully. These abovementioned tools also rely on DBs which are full of redundant, poorly characterized sequences, a fact that penalize data retrieval. The major advantages of excluding such useless sequences from a DB is that it can provide a platform to analyze the type of data that the user is more interested in. Thus, by combining the use of functional-oriented DBs and the proper methodology for genomic annotation, we created a universal tool that can potentially be utilized across different genomic sources (bacteria, archaea and metagenomes) and can annotate functional data in variety of flavors (E.C., KO, and eggNOG numbers/identifiers).

We trained our program with the genome of *E. coli* K-12, the MAG of *Ferrovum myxofaciens* S2.4, and a freshwater metagenome where Arche showed good results among all tools tested (Figs. 3-5). However, when it was run on the MAG of *F. myxofaciens* (Fig. 4), besides of having numbers of validated annotations comparable to other tools, Arche showed a higher proportion of annotations that haven’t been validated. This is also observed for other software like MicrobeAnnotator (Fig. 4) [27]. In the case of Arche, these non-validated annotations could be i) false positives, ii) positive annotations for those CDSs unannotated by other tools, iii) alternative annotations which result from the utilization of different databases, or iv) a combination of the before mentioned. The fact of the input being a MAG could also be affecting the results as the strategy used by Arche for gene calling, GeneMarkS-2, wasn’t originally intended for MAGs[30]. On the other hand, the number of false positives should be reduced through a rising in the stringency of the search parameters, like the E-value threshold and the query cover threshold which are both set by default in 10^−8^, and 70% respectively.

**Fig. 4.**
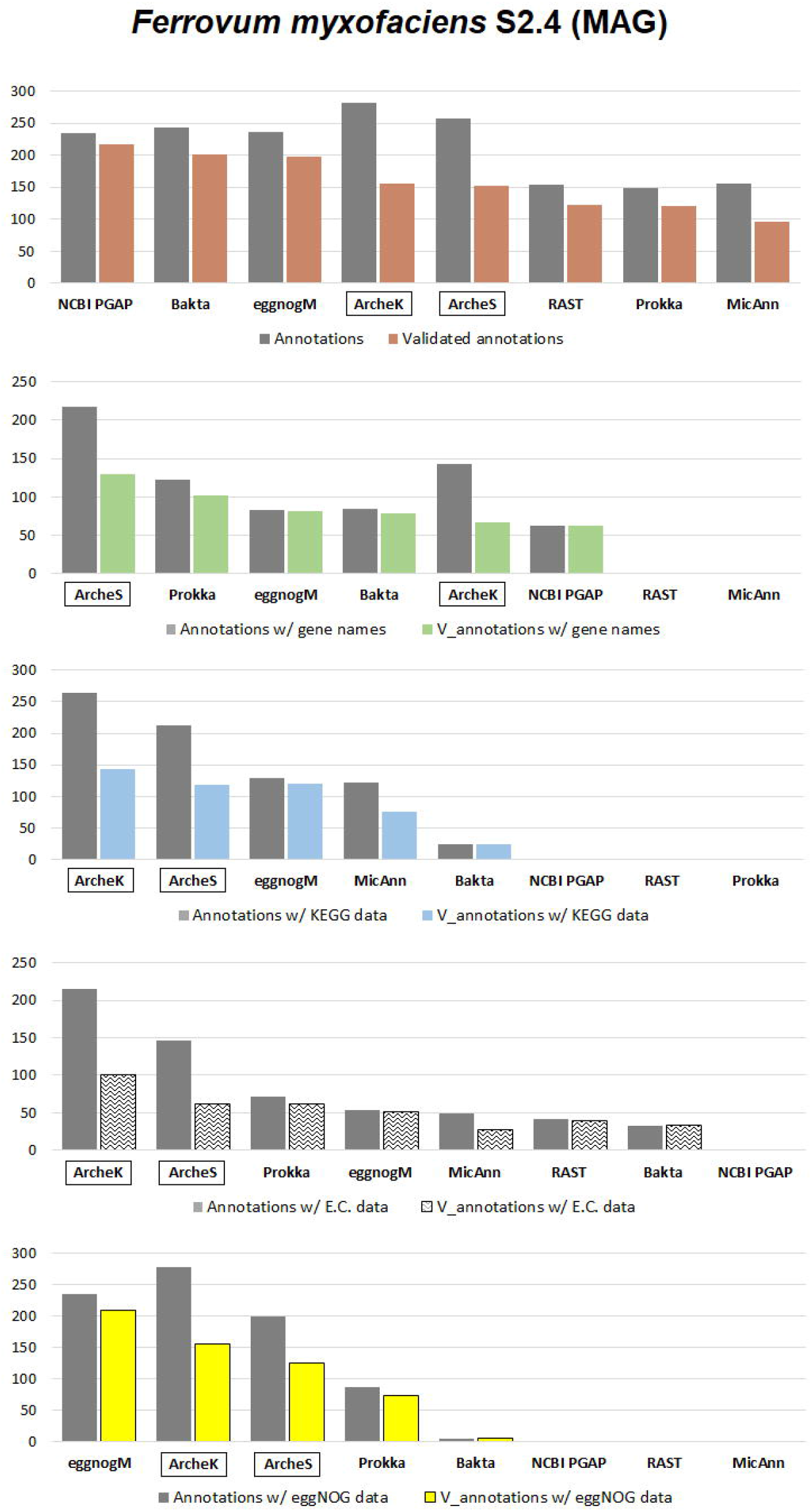
Bar plot that shows the performance of several annotation tools in the genome of the MAG *Ferrovum myxofaciens* S2.4. Abbr. V_annotations = Validated annotations

**Fig. 5.**
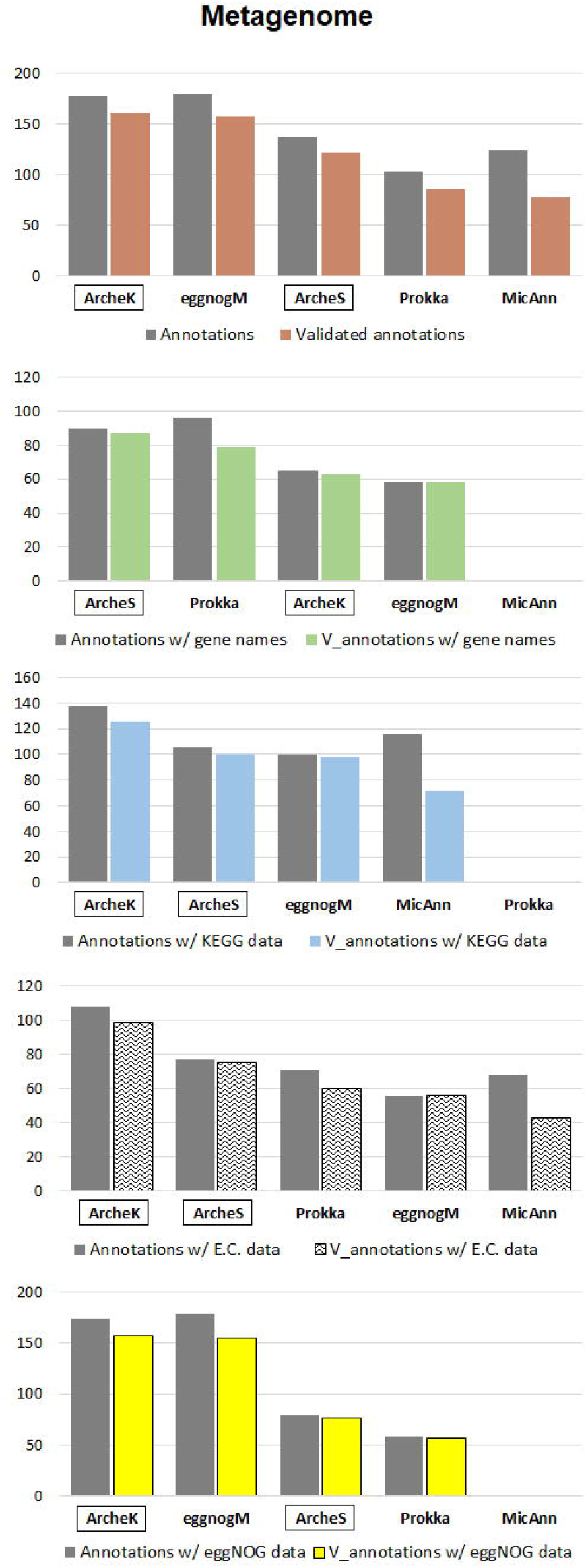
Bar plot that shows the performance of several annotation tools in a freshwater metagenome. Abbr. V_annotations = Validated annotations.

**Fig. 6.**
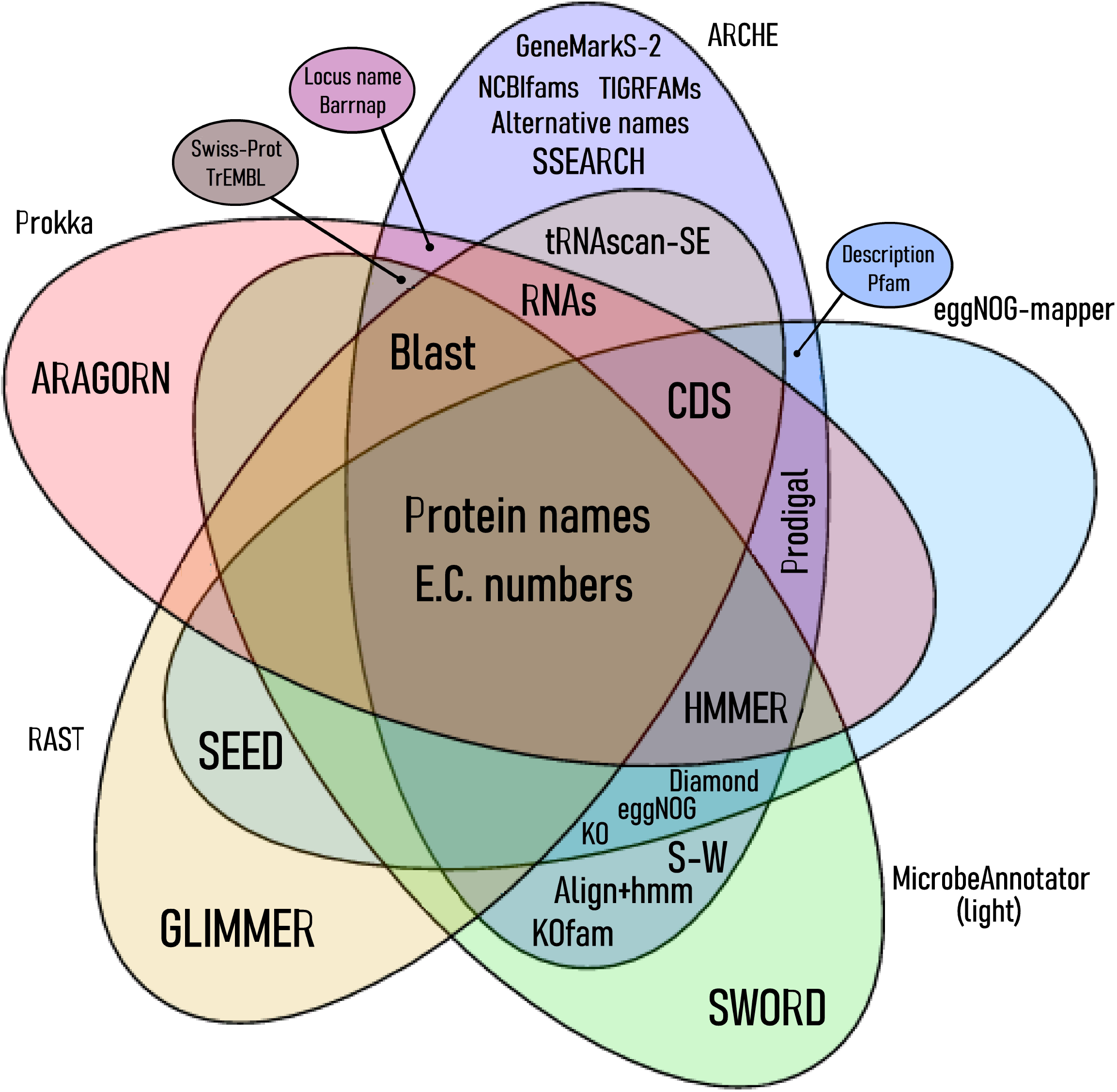
Set of shared and unshared tools and procedures of five different annotators, including Arche.

Arche was also compared with Bakta, a tool intended for bacterial genomes, which uses an alignment-free hash-based approach. Alignment-free approaches can be defined as any method of quantifying sequence similarity/dissimilarity that does not use or produce alignment (assignment of residue–residue correspondence) at any step of algorithm application[42], [43]. Far from the more traditional homology-based methods like the used by Arche, alignment-free approaches are slowly gaining popularity because they are computationally less expensive, among another advantages[42], [43]. Bakta has a better performance than Arche in the annotation of both the genome of *E. coli* K-12, and the MAG of *F. myxofaciens* S2.4. However, Bakta performed poorly in the retrieving of KEGG identifiers from *E. coli* K-12 (Fig. 2). Also KEGGs, eggNOGs, and E.C.s functional identifiers were scarce in the Bakta’s annotation of the MAG (Fig. 3).

Very recently, different annotation approaches were adopted for genomic and metagenomic analysis by our group, including the annotation of metagenomic reads through custom gene-based DBs[44], Prokka (with custom DBs) annotation of genomic contigs[45], and HMMER screening using specific gene profiles[46]. All our previous analysis lacked of a fast, standardized pipeline capable of giving rich outputs of useful metabolic data. That would require the right DBs combined with a carefully picking of all available functional codes (E.C., KO, eggNOG) that such DBs could provide. Arche overcomes the previous limitations by the implementation of NRBDs along with their associated mapping tables (Figs. 1, 2) to provide a more robust and high-throughput analysis of genomes and metagenomes.

In sum, the Arche bash-based command-line tool for genomic annotation offers an automated, competitive, and flexible workflow that can yield high-quality results for further metabolic studies.

## V. Method details

The annotation tools tested include Prokka v1.14.6, RAST v2.0, eggNOG-Mapper v2.1.3, MicrobeAnnotator light v.2.0.5, Bakta 1.9.3, and Arche v.1.0.1. All the essays were run with an E-value threshold of 10^−8^, 70% of query cover threshold, and 20 threads, using an Intel(R) Core(TM) i9-10850K computer with Ubuntu v. 24.04. Both Arche and MicrobeAnnotator tools were run with BLAST search option. A fasta file with sequences of proteins predicted by GeneMarkS-2 was provided as input for MicrobeAnnotator.

To test how comprehensive and consistent the annotations of each tool are, we annotated the genome of *E. coli* K-12 (NCBI NC_000913.3), and the freshwater metagenome (GCA_900143125.1). Then we quantified the number CDSs having attached a name denoting some functionality and functional identifiers (e. g. E.C. numbers) in the output. We considered a protein annotated if it had at least one match in any of its DBs, which would transfer both a functional name (e. g. “DNA polymerase”) and a functional ID/code/number (E.C., eggNOG and/or KO) to the queried CDS. Thus, all uninformative annotations like those labeled as “hypothetical protein”, “putative protein”, “uncharacterized protein”, “no match found”, “non supervised orthologous group”, “family of unknown function”, “domain of unknown function”, “protein of unknown function”, “domain protein”, “uncharacterized conserved protein”, or “DUF protein” with any clue of function, additional name or description will be considered as unannotated or nulls annotations. In the case of eggNOG-mapper a protein is considered annotated when the match had transferred a name that denotes some function, description or a KO identifier (e.g., K02313) to the queried CDS; in case that the annotation includes one of the uninformative labels mentioned before and doesn’t include additional functional information it will be discarded as unannotated or null annotation.

We evaluate the quality of the Arche’s annotation through a concordance analysis against other related tools in the genome of *E. coli* K12 (NCBI NC_000913.3), the metagenome-assembled genome of *Ferrovum myxofaciens* S2.4 (NZ_CP053676.1), and the freshwater metagenome (GCA_900143125.1). We measure the number of concordances in the first 250-300 CDS of the genomes, checking for the relative behavior of each tool. If the annotation made by tool has at least one correspondence by a different tool, the annotation is considered valid. We certify correspondences through the labeled names, alternative names, gene names, and functional identifiers (KO, EC, and eggNOG). More than 5250 annotations were analyzed.

## Acknowledgment

The authors acknowledge the generous financial support by PICT RAICES 2019–3216 projects. V. H. A. as a Georg Forster Experienced Researcher acknowledges the Alexander von Humboldt Foundation, which has also financially supported this project. V. H. A. is a staff researcher from the National Research Council (CONICET) in Argentina. D. A. is a postdoctoral researcher at the Center for Electron Microscopy (CIME). This manuscript has been released as a Pre-Print at bioRxiv.

