## Supplementary Figs. for "^1^Arche: An Advanced Flexible Tool for High-throuput Annotation of Functions on Microbial Contigs"

### (Supplementary figures)

| | | | | | | | | | | |
|---|---|---|---|---|---|---|---|---|---|---|
| 1 | NC_000913.3:3-98 | native | P00561 | Bifunctional aspartokinase/homoserine dehydrogenase 1 | Aspartokinase I/homoserine dehydrogenase I | thrA | 2.7.2.4 1.1.1.3 1.1.1.3 1.1.1.3 2.7.2.4 | K12524 | COG0460 | |
| 2 | NC_000913.3:337-2799 | native | P00561 | Bifunctional aspartokinase/homoserine dehydrogenase 1 | Aspartokinase I/homoserine dehydrogenase I | thrA | 2.7.2.4 1.1.1.3 1.1.1.3 1.1.1.3 2.7.2.4 | K12524 | COG0460 | |
| 3 | NC_000913.3:2801-3733 | native | P00561 | Bifunctional aspartokinase/homoserine dehydrogenase 1 | Aspartokinase I/homoserine dehydrogenase I | thrA | 2.7.2.4 1.1.1.3 1.1.1.3 1.1.1.3 2.7.2.4 | K12524 | COG0460 | |
| 4 | NC_000913.3:3734-5020 | native | P00561 | Bifunctional aspartokinase/homoserine dehydrogenase 1 | Aspartokinase I/homoserine dehydrogenase I | thrA | 2.7.2.4 1.1.1.3 1.1.1.3 1.1.1.3 2.7.2.4 | K12524 | COG0460 | |
| 5 | NC_000913.3:5234-5530 | native | P75616 | Uncharacterized protein YaaX | | yaaX | | | ENOG5032XJS | |
| 6 | NC_000913.3:5683-6459 | native | B1XBD1 | UPF0246 protein YaaA | | yaaA | | K09861 | | |
| 7 | NC_000913.3:6529-7911 | native | P00561 | Bifunctional aspartokinase/homoserine dehydrogenase 1 | Aspartokinase I/homoserine dehydrogenase I | thrA | 2.7.2.4 1.1.1.3 1.1.1.3 1.1.1.3 2.7.2.4 | K12524 | COG0460 | |
| 8 | NC_000913.3:8238-9191 | native | Q3Z606 | Transaldolase 1 | | tal1 | 2.2.1.2 2.2.1.2 2.2.1.2 | K00616 | | Transaldolase is important for the balance of metabolites in the pentose-phosphate pathway. |
| 9 | NC_000913.3:9306-9893 | native | P0AF03 | Molybdopterin adenylyltransferase | | mog | 2.7.7.75 2.7.7.75 | K03831 | COG0521 | Catalyzes the adenylation of molybdopterin as part of the biosynthesis of the molybdenum-cofactor. |
| 10 | NC_000913.3:9928-10494 | native | P0AC99 | Succinate-acetate/proton symporter SatP | Succinate-acetate transporter protein | satP | | K07034 | COG1584 | Uptake of acetate and succinate. Transport is energetically dependent on the protonmotive force (By si |
| 11 | NC_000913.3:10643-11356 | native | P75617 | UPF0174 protein YaaW | | yaaW | | | COG4735 | |
| 12 | NC_000913.3:11382-11786 | native | P28696 | UPF0412 protein YaaI | | yaaI | | | ENOG502ZXCI | |
| 13 | NC_000913.3:12163-14079 | native | Q32KA5 | Chaperone protein DnaK | HSP70 / Heat shock 70 kDa protein / Heat shock protein 70 | dnaK | | K04043 | | Acts as a chaperone. |
| 14 | NC_000913.3:14168-15298 | native | P08622 | Chaperone protein DnaJ | HSP40 / Heat shock protein J | dnaJ | | K03686 | COG0484 | Interacts with DnaK and GrpE to disassemble a protein complex at the origins of replication of phage l |
| 15 | NC_000913.3:15445-16557 | atypical | P0CF92 | Putative transposase InsL for insertion sequence element IS186 | | insL2 | | | | |
| 16 | NC_000913.3:16751-16903 | atypical | P0ACG5 | Protein HokC | | hokC | | K18919 | | Toxic component of a type I toxin-antitoxin (TA) system (By similarity). When overexpressed kills cell |
| 17 | NC_000913.3:17489-18655 | native | P13738 | Na(+)/H(+) antiporter NhaA | Sodium/proton antiporter NhaA | nhaA | | K03313 | COG3004 | Na(+)/H(+) antiporter that extrudes sodium in exchange for external protons. Catalyzes the exchange of |
| 18 | NC_000913.3:18715-19620 | native | P0A9G2 | Transcriptional activator protein NhaR | Na(+)/H(+) antiporter regulatory protein | nhaR | | K03717 | COG0583 | Plays a role in the positive regulation of NhaA. |
| 19 | NC_000913.3:19811-20188 | native | P0CF29 | Insertion element IS1 6 protein InsB | IS1e | insB6 | | K07480 | | |
| 20 | NC_000913.3:20233-20508 | native | P0CF07 | Insertion element IS1 1 protein InsA | IS1a | insA1 | | | COG3677 | |
| 21 | NC_000913.3:20815-21078 | atypical | B5YYB6 | 30S ribosomal protein S20 | | rpsT | | K02968 | | Binds directly to 16S ribosomal RNA. |
| 22 | NC_000913.3:21407-22348 | native | P0AG41 | Bifunctional riboflavin kinase/FMN adenylyltransferase | Riboflavin biosynthesis protein RibF / Flavokinase / FAD pyrophosphorylase | ribF | 2.7.1.26 2.7.7.2 2.7.1.26 2.7.7.2 2.7.1.26 2.7.7.2 | K11753 | COG0196 | Catalyzes the phosphorylation of riboflavin to FMN followed by the adenylation of FMN to FAD. |
| 23 | NC_000913.3:22391-25207 | native | B6HZ21 | Isoleucine--tRNA ligase | Isoleucyl-tRNA synthetase | ileS | 6.1.1.5 6.1.1.5 6.1.1.5 | K01870 | | Catalyzes the attachment of isoleucine to tRNA(Ile). As IleRS can inadvertently accommodate and proces |
| 24 | NC_000913.3:25207-25701 | native | Q3Z5Y3 | Lipoprotein signal peptidase | Prolipoprotein signal peptidase / Signal peptidase II | lspA | 3.4.23.36 3.4.23.36 3.4.23.36 | K03101 | | This protein specifically catalyzes the removal of signal peptides from prolipoproteins. |
| 25 | NC_000913.3:25826-26275 | native | P0AEM0 | FKBP-type 16 kDa peptidyl-prolyl cis-trans isomerase | Rotamase | fkpB | 5.2.1.8 5.2.1.8 | K03774 | COG1047 | PPIases accelerate the folding of proteins. Substrate specificity investigated with 'Suc-Ala-Xaa-Pro-P |
| 26 | NC_000913.3:26277-27227 | native | P0AG41 | Bifunctional riboflavin kinase/FMN adenylyltransferase | Riboflavin biosynthesis protein RibF / Flavokinase / FAD pyrophosphorylase | ribF | 2.7.1.26 2.7.7.2 2.7.1.26 2.7.7.2 2.7.1.26 2.7.7.2 | K11753 | COG0196 | Catalyzes the phosphorylation of riboflavin to FMN followed by the adenylation of FMN to FAD. |
| 27 | NC_000913.3:27293-28207 | native | C4ZPV6 | Non-specific ribonucleoside hydrolase RihC | Purine/pyrimidine ribonucleoside hydrolase | rihC | 3.2.-.- 3.2.-.- | K12700 | | Hydrolyzes both purine and pyrimidine ribonucleosides with a broad-substrate specificity. |
| 28 | NC_000913.3:28374-29195 | native | B7N7Q5 | 4-hydroxy-tetrahydrodipicolinate reductase | | dapB | 1.17.1.8 1.17.1.8 1.17.1.8 1.17.1.8 | K00215 | | Catalyzes the conversion of 4-hydroxy-tetrahydrodipicolinate (HTPA) to tetrahydrodipicolinate. |
| 29 | NC_000913.3:29651-30799 | native | P0A6F1 | Carbamoyl-phosphate synthase small chain | Carbamoyl-phosphate synthetase glutamine chain | carA | 6.3.5.5 6.3.5.5 | K01956 | COG0505 | |
| 30 | NC_000913.3:30817-34038 | native | P00968 | Carbamoyl-phosphate synthase large chain | Carbamoyl-phosphate synthetase ammonia chain | carB | 6.3.5.5 6.3.5.5 6.3.5.5 | K01955 | COG0458 | |
| 31 | NC_000913.3:34300-34695 | native | P0AE58 | Transcriptional activatory protein CaiF | | caiF | | K08277 | ENOG5030W1N | |
| 32 | NC_000913.3:34781-35371 | native | B1XBG0 | Carnitine operon protein CaiE | | caiE | | K08279 | | Overproduction of CaiE stimulates the activity of CaiB and CaiD. |
| 33 | NC_000913.3:35377-36162 | native | P31551 | Carnitinyl-CoA dehydratase | Crotonobetainyl-CoA hydratase | caiD | 4.2.1.149 4.2.1.149 | K08299 | COG1024 | Catalyzes the reversible dehydration of L-carnitinyl-CoA to crotonobetainyl-CoA. |
| 34 | NC_000913.3:36271-37839 | native | B1XBG2 | Crotonobetaine/carnitine--CoA ligase | | caiC | 6.2.1.48 6.2.1.48 6.2.1.48 6.2.1.48 6.2.1.48 | K02182 | | Catalyzes the transfer of CoA to carnitine, generating the initial carnitinyl-CoA needed for the CaiB |
| 35 | NC_000913.3:37898-39115 | native | B1IRD8 | L-carnitine CoA-transferase | Crotonobetainyl-CoA:carnitine CoA-transferase | caiB | 2.8.3.21 2.8.3.21 2.8.3.21 2.8.3.21 | K08298 | | Catalyzes the reversible transfer of the CoA moiety from gamma-butyrobetainyl-CoA to L-carnitine to ge |
| 36 | NC_000913.3:39244-40386 | native | Q3Z5Y3 | Lipoprotein signal peptidase | Prolipoprotein signal peptidase / Signal peptidase II | lspA | 3.4.23.36 3.4.23.36 3.4.23.36 | K03101 | | This protein specifically catalyzes the removal of signal peptides from prolipoproteins. |
| 37 | NC_000913.3:40417-41931 | native | C4ZPW6 | L-carnitine/gamma-butyrobetaine antiporter | | caiT | | K05245 | | Catalyzes the exchange of L-carnitine for gamma-butyrobetaine and related betaines. |
| 38 | NC_000913.3:42403-43173 | native | B1XBG6 | Protein FixA | | fixA | | K03521 | | Required for anaerobic carnitine reduction. May bring reductant to CaiA. |
| 39 | NC_000913.3:43188-44129 | native | P00547 | Homoserine kinase | | thrB | 2.7.1.39 2.7.1.39 | K00872 | COG0083 | Catalyzes the ATP-dependent phosphorylation of L-homoserine to L-homoserine phosphate. Is also able to |
| 40 | NC_000913.3:44180-45466 | native | P68645 | Protein FixC | | fixC | | K00313 | COG0644 | Could be part of an electron transfer system required for anaerobic carnitine reduction. |
| 41 | NC_000913.3:45463-45750 | native | P68646 | Ferredoxin-like protein FixX | | fixX | | K03855 | COG2440 | |
| 42 | NC_000913.3:45807-47138 | native | P31679 | Putative metabolite transport protein YaaU | | yaaU | | K08368 | COG2271 | |
| 43 | NC_000913.3:47246-47776 | native | B1XC47 | Glutathione-regulated potassium-efflux system ancillary protei | Quinone oxidoreductase KefF | kefF | 1.6.5.2 1.6.5.2 1.6.5.2 1.6.5.2 | K11746 | | Regulatory subunit of a potassium efflux system that confers protection against electrophiles. Require |
| 44 | NC_000913.3:47769-49631 | native | P03819 | Glutathione-regulated potassium-efflux system protein KefC | K(+)/H(+) antiporter | kefC | | K11745 | COG0475 | Pore-forming subunit of a potassium efflux system that confers protection against electrophiles. Catal |

**Supplementary Figure 1.** Snippet from an example []_omic_table.tbl file, when applied the command *column -ts "|" [example]_omic_table.tbl | less -S*

| | A | B | C | D | E | F | G | H | I | J | K |
|---|---|---|---|---|---|---|---|---|---|---|---|
| 1 | 1 | NC_000913.3:3-98 | native | P00561 | Bifunctional aspartokinase/homoserine dehydrogenase 1 | Aspartokinase I/homoserine dehydrogenase I | thrA | 2.7.2.4 1.1.1.3 1.1.1.3 1.1.1.3 2.7.2.4 | K12524 | COG0460 | |
| 2 | 2 | NC_000913.3:337-2799 | native | P00561 | Bifunctional aspartokinase/homoserine dehydrogenase 1 | Aspartokinase I/homoserine dehydrogenase I | thrA | 2.7.2.4 1.1.1.3 1.1.1.3 1.1.1.3 2.7.2.4 | K12524 | COG0460 | |
| 3 | 3 | NC_000913.3:2801-3733 | native | P00561 | Bifunctional aspartokinase/homoserine dehydrogenase 1 | Aspartokinase I/homoserine dehydrogenase I | thrA | 2.7.2.4 1.1.1.3 1.1.1.3 1.1.1.3 2.7.2.4 | K12524 | COG0460 | |
| 4 | 4 | NC_000913.3:3734-5020 | native | P00561 | Bifunctional aspartokinase/homoserine dehydrogenase 1 | Aspartokinase I/homoserine dehydrogenase I | thrA | 2.7.2.4 1.1.1.3 1.1.1.3 1.1.1.3 2.7.2.4 | K12524 | COG0460 | |
| 5 | 5 | NC_000913.3:5234-5530 | native | P75616 | Uncharacterized protein YaaX | | yaaX | | | ENOG5032XJS | |
| 6 | 6 | NC_000913.3:5683-6459 | native | B1XBD1 | UPF0246 protein YaaA | | yaaA | | K09861 | | |
| 7 | 7 | NC_000913.3:6529-7911 | native | P00561 | Bifunctional aspartokinase/homoserine dehydrogenase 1 | Aspartokinase I/homoserine dehydrogenase I | thrA | 2.7.2.4 1.1.1.3 1.1.1.3 1.1.1.3 2.7.2.4 | K12524 | COG0460 | |
| 8 | 8 | NC_000913.3:8238-9191 | native | Q3Z606 | Transaldolase 1 | | tal1 | 2.2.1.2 2.2.1.2 2.2.1.2 | K00616 | | Transaldolase is important for the balance of metabolites in the pentose-phosphate pathway. |
| 9 | 9 | NC_000913.3:9306-9893 | native | P0AF03 | Molybdopterin adenylyltransferase | | mog | 2.7.7.75 2.7.7.75 | K03831 | COG0521 | Catalyzes the adenylation of molybdopterin as part of the biosynthesis of the molybdenum-cofactor. |
| 10 | 10 | NC_000913.3:9928-10494 | native | P0AC99 | Succinate-acetate/proton symporter SatP | Succinate-acetate transporter protein | satP | | K07034 | COG1584 | Uptake of acetate and succinate. Transport is energetically dependent on the protonmotive force (By similarity). |
| 11 | 11 | NC_000913.3:10643-11356 | native | P75617 | UPF0174 protein YaaW | | yaaW | | | COG4735 | |
| 12 | 12 | NC_000913.3:11382-11786 | native | P28696 | UPF0412 protein YaaI | | yaaI | | | ENOG502ZXCI | |
| 13 | 13 | NC_000913.3:12163-14079 | native | Q32KA5 | Chaperone protein DnaK | HSP70 / Heat shock 70 kDa protein / Heat shock protein 70 | dnaK | | K04043 | | Acts as a chaperone. |
| 14 | 14 | NC_000913.3:14168-15298 | native | P08622 | Chaperone protein DnaJ | HSP40 / Heat shock protein J | dnaJ | | K03686 | COG0484 | Interacts with DnaK and GrpE to disassemble a protein complex at the origins of replication of phage lambda and several plasmids. Par |
| 15 | 15 | NC_000913.3:15445-16557 | atypical | P0CF92 | Putative transposase InsL for insertion sequence element IS186B | | insL2 | | | | |
| 16 | 16 | NC_000913.3:16751-16903 | atypical | P0ACG5 | Protein HokC | | hokC | | K18919 | | Toxic component of a type I toxin-antitoxin (TA) system (By similarity). When overexpressed kills cells within minutes; causes collaps |
| 17 | 17 | NC_000913.3:17489-18655 | native | P13738 | Na(+)/H(+) antiporter NhaA | Sodium/proton antiporter NhaA | nhaA | | K03313 | COG3004 | Na(+)/H(+) antiporter that extrudes sodium in exchange for external protons. Catalyzes the exchange of 2 H(+) per Na(+). Can mediate |
| 18 | 18 | NC_000913.3:18715-19620 | native | P0A9G2 | Transcriptional activator protein NhaR | Na(+)/H(+) antiporter regulatory protein | nhaR | | K03717 | COG0583 | Plays a role in the positive regulation of NhaA. |
| 19 | 19 | NC_000913.3:19811-20188 | native | P0CF29 | Insertion element IS1 6 protein InsB | IS1e | insB6 | | K07480 | | |
| 20 | 20 | NC_000913.3:20233-20508 | native | P0CF07 | Insertion element IS1 1 protein InsA | IS1a | insA1 | | | COG3677 | |
| 21 | 21 | NC_000913.3:20815-21078 | atypical | B5YYB6 | 30S ribosomal protein S20 | | rpsT | | K02968 | | Binds directly to 16S ribosomal RNA. |
| 22 | 22 | NC_000913.3:21407-22348 | native | P0AG41 | Bifunctional riboflavin kinase/FMN adenylyltransferase | Riboflavin biosynthesis protein RibF / Flavokinase / FAD pyrophosphorylase / FAD s | ribF | 2.7.1.26 2.7.7.2 2.7.1.26 2.7.7.2 2.7.1.26 2.7.7.2 | K11753 | COG0196 | Catalyzes the phosphorylation of riboflavin to FMN followed by the adenylation of FMN to FAD. |
| 23 | 23 | NC_000913.3:22391-25207 | native | B6HZ21 | Isoleucine--tRNA ligase | Isoleucyl-tRNA synthetase | ileS | 6.1.1.5 6.1.1.5 6.1.1.5 | K01870 | | Catalyzes the attachment of isoleucine to tRNA(Ile). As IleRS can inadvertently accommodate and process structurally similar amino ac |
| 24 | 24 | NC_000913.3:25207-25701 | native | Q3Z5Y3 | Lipoprotein signal peptidase | Prolipoprotein signal peptidase / Signal peptidase II | lspA | 3.4.23.36 3.4.23.36 3.4.23.36 | K03101 | | This protein specifically catalyzes the removal of signal peptides from prolipoproteins. |
| 25 | 25 | NC_000913.3:25826-26275 | native | P0AEM0 | FKBP-type 16 kDa peptidyl-prolyl cis-trans isomerase | Rotamase | fkpB | 5.2.1.8 5.2.1.8 | K03774 | COG1047 | PPIases accelerate the folding of proteins. Substrate specificity investigated with 'Suc-Ala-Xaa-Pro-Phe-4-nitroanilide' where Xaa is th |
| 26 | 26 | NC_000913.3:26277-27227 | native | P0AG41 | Bifunctional riboflavin kinase/FMN adenylyltransferase | Riboflavin biosynthesis protein RibF / Flavokinase / FAD pyrophosphorylase / FAD s | ribF | 2.7.1.26 2.7.7.2 2.7.1.26 2.7.7.2 2.7.1.26 2.7.7.2 | K11753 | COG0196 | Catalyzes the phosphorylation of riboflavin to FMN followed by the adenylation of FMN to FAD. |
| 27 | 27 | NC_000913.3:27293-28207 | native | C4ZPV6 | Non-specific ribonucleoside hydrolase RihC | Purine/pyrimidine ribonucleoside hydrolase | rihC | 3.2.-.- 3.2.-.- | K12700 | | Hydrolyzes both purine and pyrimidine ribonucleosides with a broad-substrate specificity. |
| 28 | 28 | NC_000913.3:28374-29195 | native | B7N7Q5 | 4-hydroxy-tetrahydrodipicolinate reductase | | dapB | 1.17.1.8 1.17.1.8 1.17.1.8 1.17.1.8 | K00215 | | Catalyzes the conversion of 4-hydroxy-tetrahydrodipicolinate (HTPA) to tetrahydrodipicolinate. |
| 29 | 29 | NC_000913.3:29651-30799 | native | P0A6F1 | Carbamoyl-phosphate synthase small chain | Carbamoyl-phosphate synthetase glutamine chain | carA | 6.3.5.5 6.3.5.5 | K01956 | COG0505 | |
| 30 | 30 | NC_000913.3:30817-34038 | native | P00968 | Carbamoyl-phosphate synthase large chain | Carbamoyl-phosphate synthetase ammonia chain | carB | 6.3.5.5 6.3.5.5 6.3.5.5 | K01955 | COG0458 | |
| 31 | 31 | NC_000913.3:34300-34695 | native | P0AE58 | Transcriptional activatory protein CaiF | | caiF | | K08277 | ENOG5030W1N | |
| 32 | 32 | NC_000913.3:34781-35371 | native | B1XBG0 | Carnitine operon protein CaiE | | caiE | | K08279 | | Overproduction of CaiE stimulates the activity of CaiB and CaiD. |
| 33 | 33 | NC_000913.3:35377-36162 | native | P31551 | Carnitinyl-CoA dehydratase | Crotonobetainyl-CoA hydratase | caiD | 4.2.1.149 4.2.1.149 | K08299 | COG1024 | Catalyzes the reversible dehydration of L-carnitinyl-CoA to crotonobetainyl-CoA. |
| 34 | 34 | NC_000913.3:36271-37839 | native | B1XBG2 | Crotonobetaine/carnitine--CoA ligase | | caiC | 6.2.1.48 6.2.1.48 6.2.1.48 6.2.1.48 6.2.1.48 | K02182 | | Catalyzes the transfer of CoA to carnitine, generating the initial carnitinyl-CoA needed for the CaiB reaction cycle. Also has activity tov |
| 35 | 35 | NC_000913.3:37898-39115 | native | B1IRD8 | L-carnitine CoA-transferase | Crotonobetainyl-CoA:carnitine CoA-transferase | caiB | 2.8.3.21 2.8.3.21 2.8.3.21 2.8.3.21 | K08298 | | Catalyzes the reversible transfer of the CoA moiety from gamma-butyrobetainyl-CoA to L-carnitine to generate L-carnitinyl-CoA and g |
| 36 | 36 | NC_000913.3:39244-40386 | native | Q3Z5Y3 | Lipoprotein signal peptidase | Prolipoprotein signal peptidase / Signal peptidase II | lspA | 3.4.23.36 3.4.23.36 3.4.23.36 | K03101 | | This protein specifically catalyzes the removal of signal peptides from prolipoproteins. |
| 37 | 37 | NC_000913.3:40417-41931 | native | C4ZPW6 | L-carnitine/gamma-butyrobetaine antiporter | | caiT | | K05245 | | Catalyzes the exchange of L-carnitine for gamma-butyrobetaine and related betaines. |
| 38 | 38 | NC_000913.3:42403-43173 | native | B1XBG6 | Protein FixA | | fixA | | K03521 | | Required for anaerobic carnitine reduction. May bring reductant to CaiA. |

**Supplementary Figure 2.** Snippet from an example []_omic_table.tsv file

```
Runing Arche with the following parameters:
input = ECOLIK12.fasta
kingdom = bacteria
mode = standard
cpus = 20
e-value = 1e-08
memory = 40 GB
bypass RNA prediction = no
gene prediction = genemark
query coverage = 70 %
verbose mode = off

Results:
15 rRNAs
89 tRNAs
4269 putative protein(s) (3749 native and 520 atypical)
4214 standard (SwissProt) match(es)
1 KEGG (from HMMs)
0 match(es) with NFAM
1 match(es) with TIGRFAM
2 match(es) with PFAM
51 hypotetical protein(s)
Total duration of the process: 378 seconds.
~
~
```

**Supplementary Figure 3.** Snippet from an example arche_report file
